# Distinct Neural Signatures of Auditory Processing in Contact versus Non-contact Sports Athletes

**DOI:** 10.64898/2026.02.13.705730

**Authors:** Jessica R. Andrew, Ewan Dean, Andrew Thomas, Christopher J. Plack, Christopher J. Gaffney, Helen E. Nuttall

**Affiliations:** Department of Psychology, Lancaster University, Lancaster, LA1 4AT, UK; Lancaster Medical School, Sir John Fisher Drive, Lancaster University, Lancaster, LA1 4AT, UK; Brainbox Ltd, 6 Museum Place, Cardiff, CF10 3BG, UK; Manchester Centre for Audiology and Deafness, School of Health Sciences, The University of Manchester, Manchester, M13 9PL UK

**Author notes:** Correspondence to:* Dr. Helen Nuttall.

**Keywords:** Brain, Concussion, Sub-concussion, Contact sports, Auditory System

## Abstract

Repetitive sub-concussive head impacts are emerging as one of the most urgent and overlooked challenges in neurotrauma. Despite growing evidence of their neurological consequences, there is no validated objective biomarker for early and reliable detection. Cortical (N100) and subcortical (frequency following responses) to a speech syllable presented in (1) quiet and (2) six-talker background noise listening conditions were assessed using EEG in 60 tier-2 athletes (30 contact, 30 non-contact; age-, sex-, height-, body mass- and BMI-matched). Reduced cortical N100 amplitudes in contact athletes were confirmed by a significant group effect (*F*(1,54) = 9.16, *p* = .004), indicating early auditory cortical dysfunction as a measurable biomarker of sub-concussive exposure. Contact athletes also exhibited subtle hearing deficits and impaired self-reported speech perception, linking neural changes to real-world communication deficits. These findings were not related to cortical response amplitudes, suggesting that peripheral and cortical changes may occur independently following repetitive head impacts. Response timing and subcortical encoding were unaffected under both listening conditions. Our findings establish a selective auditory cortical vulnerability to repeated sub-concussive head impact exposure, providing the basis of an objective EEG-based monitoring tool that could help support athlete brain health and safety, and inform future research in contact sports.

## Introduction

Head injuries in sport are a topic of concern, due to the consequences on physical health, cognitive function, and long-term brain health^[1]^. In recent years, growing attention has turned to the more subtle head impacts known as “sub-concussive” or “non-concussive” head impacts^[2]^. Although there remains no universally agreed terminology, both terms refer to repetitive head impacts that do not produce overt concussion symptoms but may still have cumulative, long-term effects^[3]^. Heading in football exemplifies such impacts, as the brain moves rapidly inside the skull but this does not result in noticeable acute symptoms^[4]^. Emerging evidence links the repetitive nature of these head impacts, accumulated over years of play, to alterations in brain structure and function and an increased risk of neurodegenerative disease, particularly with early and prolonged exposure^[5]^. Unlike concussions which result in recognisable symptoms, sub-concussive head impacts often go unnoticed and untreated^[6]^.

Emerging research demonstrates an association between former contact sport players who have been exposed to repetitive sub-concussive exposure and an increased risk of both chronic traumatic encephalopathy (CTE), a progressive neurodegenerative disease^[7]^, and cognitive decline^[8]^ when compared with matched controls^[9]^. CTE is characterised by the perivascular accumulation of hyperphosphorylated tau in neurons and astrocytes at the depths of cortical sulci, often accompanied in more advanced stages by widespread white matter degeneration, ventricular enlargement, and cortical and medial temporal lobe atrophy, distinguishing it from other primary tauopathies such Alzheimer’s diseases^[10–12]^. While long-term consequences are of significant concern, further research into the immediate and short-term effects of such head impacts is needed to gain a more comprehensive understanding of potential associations between these head impacts and neurodegenerative diseases^23^.

Auditory sensory deficits are a potential symptom of head injury, but their prevalence and clinical significance remain incompletely understood^[13]^. Although the present study focuses on repetitive sub-concussive exposure, related work on concussive head impacts indicate that contact sport athletes frequently experience auditory-related challenges, particularly when processing speech in background noise^[14,15]^. Early research suggested that this was due to damage to peripheral auditory function, but perspectives have recently shifted towards these symptoms arising due to central auditory processing deficits^[16]^. The central auditory system includes ascending and descending neural pathways extending from the brainstem through the thalamus to primary and secondary auditory cortices within the superior temporal gyrus^[17]^. The proximity of these cortical regions to the lateral sulcus may render them vulnerable to the shear and rotational forces that occur during repetitive sub-concussive impacts^[18]^. Such trauma could disrupt neural connectivity and synaptic communication between cortical and subcortical auditory networks, impairing accurate sound processing, a difficulty frequently reported by contact sport athletes following head impact^[19]^.

Traditional neuroimaging techniques, such as Computed Tomography and Magnetic Resonance Imaging scans, often fail to detect subtle changes following sub-concussive impacts^[20]^. Electrophysiological methods such as electroencephalogram (EEG) have demonstrated greater sensitivity in identifying functional disruptions following repetitive sub-concussive head impacts, especially when identifying deficits within auditory processing pathways^[21,22]^. Recent EEG studies have provided evidence for cortical alterations after repetitive sub-concussive exposure^[23]^. Fickling et al., (2021) investigated junior ice hockey players across a competitive season and reported a significant reduction in N100 amplitude using a passive listening paradigm, suggesting disrupted cortical auditory processing following repetitive sub-concussive head impacts^[24]^. The frequency-following response (FFR), an EEG-based measure thought to reflect the subcortical encoding of the rapid temporal characteristics of sounds, including speech, provides further insight into neural vulnerability. Athletes in high-contact sports exhibit degraded FFRs compared with non-contact controls, indicating altered brainstem processing likely resulting from cumulative sub-concussive exposure^[25,26]^.

Although cortical and subcortical auditory changes have each been reported after repetitive sub-concussive exposure, these levels have not been assessed together, nor under demanding, ecologically relevant listening conditions. To address this gap, the aim of the present study was to determine whether sub-concussive head impacts in invasion-based contact sports are associated with measurable alterations in both cortical (N100) and subcortical (FFR) responses during ecologically relevant auditory tasks, and determine which response is maximally sensitive and could serve as a future early-stage biomarker of brain injury.

We tested the following primary hypotheses:

- H1 - In the quiet condition, contact sport athletes exposed to repeated sub-concussive head impacts will show a reduced amplitude in the F0 component of FFR, compared to non-contact athletes
- H2 - Contact sport athletes exposed to repeated sub-concussive head impacts will show a larger decrease in F0 amplitude from the quiet to speech-in-noise condition than non-contact athletes, indicating a group x condition interaction.
- H3 - In the quiet condition, contact sport athletes exposed to repeated sub-concussive head impacts will show a reduced N100 amplitude, compared to non-contact athletes.
- H4 – Contact sport athletes exposed to repeated sub-concussive head impacts will show a larger decrease in N100 amplitude from the quiet to speech-in-noise condition than non-contact athletes, indicating a group x condition interaction.

## Results

### Sub-concussive Head Impacts Are Associated with Cortical Auditory Dysfunction

To determine whether sub-concussive injury manifested in cortical injury, we measured cortical response amplitudes between contact and non-contact athletes across quiet and Speech-in-Noise conditions. Figure 1 illustrates robust group-level differences in cortical auditory function. In the quiet listening condition, contact athletes show significantly smaller (less negative) N100 amplitudes than non-contact athletes, revealing a cortical neural distinction associated with repetitive sub-concussive head impacts.

**Figure 1.**
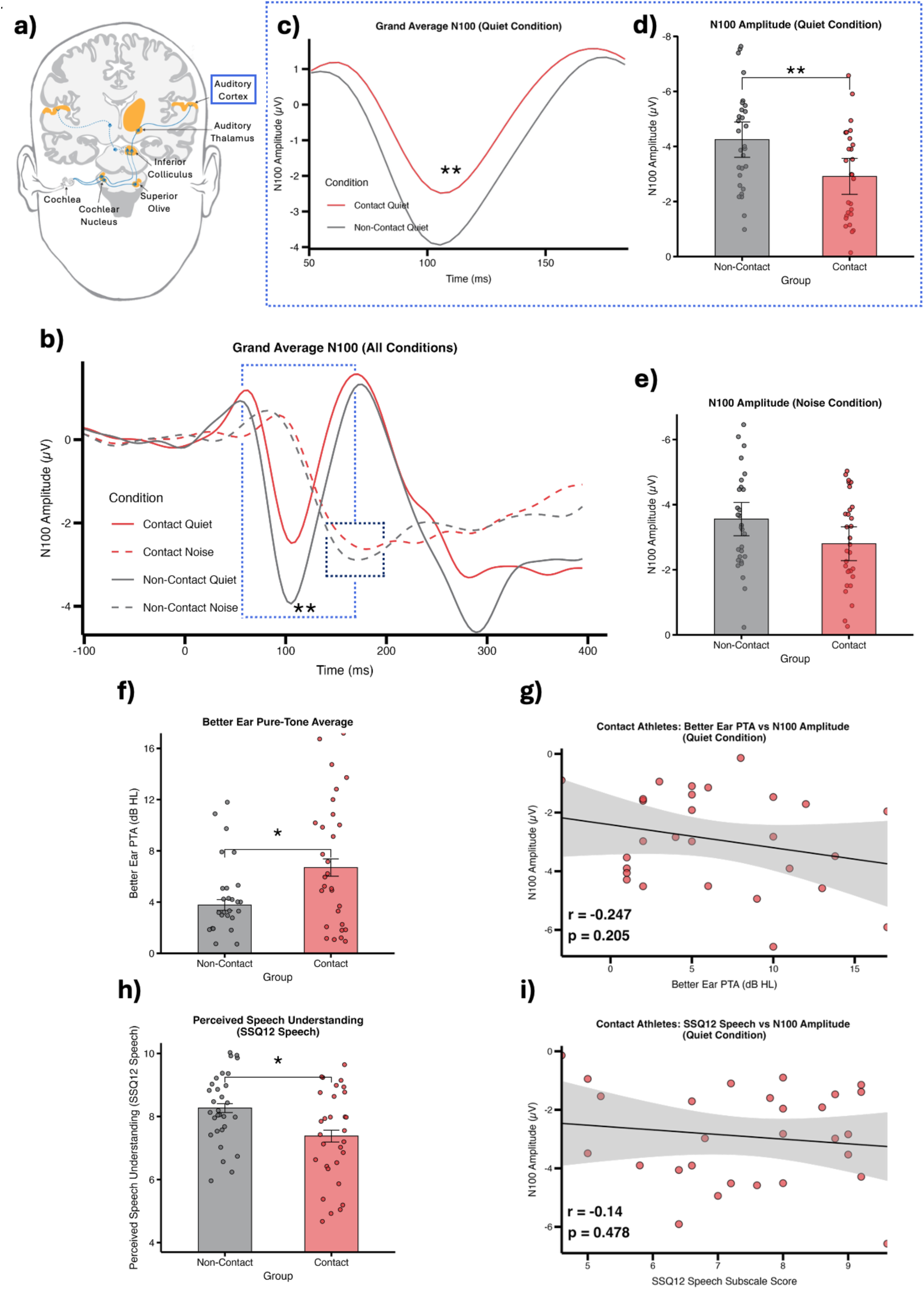
Sub-concussive head impacts are associated with cortical auditory dysfunction and reduced real-world communication ability. (a) Schematic of the human auditory pathway from cochlea to auditory cortex, highlighting regions thought to be responsible for the N100 response measured with EEG (Image adapted from Swords and colleagues^[27]^). (b) Grand-average waveforms (-100ms – 400ms) for contact and non-contact athletes across quiet and speech-in-noise (noise) listening conditions, with dashed boxes indicating N100 time window (royal blue; quiet listening condition; black; noise listening condition) and the subsequent analysis epoch of interest showing a reduced cortical response in the contact group around 100 ms. (c) Grand-average N100 in the quiet condition, illustrating markedly reduced (less negative) N100 amplitude in contact versus non-contact athletes. (d) N100 amplitude in the quiet listening condition shows significantly smaller responses in contact athletes than non-contact athletes, consistent with early-stage cortical auditory dysfunction following repetitive head impacts. (e) In the Noise condition, there was no significant difference in N100 amplitude between contact and non-contact athletes, indicating that group-level cortical responses converge when speech is presented in background noise. both groups show a less negative and delayed N100 in noise, consistent with the increased neural demands of processing speech under degraded listening conditions. Hearing thresholds and N100. (f) Contact athletes show significantly elevated better-ear pure-tone thresholds compared with non-contact athletes, indicating subtly worse hearing in the contact group relative to the non-contact athletes. (g) Within contact athletes, better-ear PTA is not significantly associated with N100 amplitude in quiet, suggesting that reduced cortical responses are not simply explained by peripheral hearing status. Real-world speech understanding. (h) Contact athletes report significantly poorer perceived speech understanding on the SSQ12 Speech subscale than non-contact peers, (i) within contact athletes, SSQ12 Speech scores are not significantly associated with N100 amplitude in quiet, indicating that cortical N100 amplitude differences arise independently of both peripheral hearing status and self-reported speech understanding and may therefore represent a selective cortical biomarker of exposure to repetitive sub-concussive head impacts.

A 2 × 2 mixed analysis of variance (ANOVA) assessing Group (Contact vs. Non-Contact) and Condition (Quiet vs. Noise) revealed a significant main effect of Group on N100 amplitude (F(1, 55) = 9.86, *p* = .003, η_p_² = .152), with contact athletes indicating smaller (less negative) amplitudes (*M* = -2.83 μV, *SD* = 1.48) than non-contact athletes (*M* = -3.83 μV, *SD* = 1.66). Applying the Bonferroni-corrected significance threshold (α = .0125), this effect remained statistically significant. The main effect of Condition was not significant (F(1, 55) = 4.71, *p* = .034, η_p_² = .079). The Group × Condition interaction was also not significant (F(1, 55) = 2.95, *p* = .158, η_p_² = .036).

### Cortical Auditory Responsiveness is Selectively Disrupted During Quiet Listening Following Repetitive Sub-Concussive Head Impacts

Planned pairwise comparisons were conducted to compare non-contact and contact athletes within each listening condition. Using the Bonferroni-corrected alpha threshold of *p* < .025. The group difference in the Quiet condition was statistically significant, t(55) = −3.10, *p* = .003, with contact athletes showing smaller N100 amplitudes than non-contact athletes. In the noise condition, the group difference did not reach significance, t(55) = −2.11, *p* = .039. These data indicate that sub-concussive head impacts from contact sports are associated with cortical auditory dysfunction, whereby reduced N100 amplitude may indicate reduced neural synchronisation or reduced neural firing.

### Cortical Dysfunction Occurs Independently of Peripheral Hearing and is Associated with Deficits in Speech Perception

Contact athletes also exhibited worse peripheral hearing in their better ear compared to non-contact athletes (t(55) = −3.694, *p* = .0004, Fig. 1f), yet within the contact group, better ear PTA thresholds showed no association with N100 amplitudes in quiet (r = -0.247, *p* = .205, Fig. 1g). Contact athletes further reported poorer real-world speech understanding on the SSQ12 Speech Subscale (t(55) = 2.655, *p* = 0.101, Fig. 1h), and within the contact group, SSQ12 Speech scores were not significantly associated with N100 amplitudes in quiet (r = -0.140, *p* = .478, Fig. 1i) indicating that the cortical N100 differences observed in contact athletes arise independently of both peripheral hearing status and self-reported speech understanding, and may therefore reflect a selective cortical biomarker of repeated sub-concussive head impacts.

### Cortical N100 Timing Preserved Despite Amplitude Deficits

To examine effects on cortical response timing, a 2 x 2 ANOVA assessed Group (Contact vs. Non-Contact) and Condition (Quiet vs. Noise) on N100 latency. A significant main effect of Condition confirmed longer latencies in the noise condition versus the quiet condition for both athlete groups, F(1, 58) = 507.59, *p* < .001, η_p_² = .897. No significant effects of Group, F(1, 58) = 0.02, *p* = .881, or Group × Condition interaction, F(1, 58) = 1.21, *p* = .277, were observed.

Pairwise comparisons showed that both groups exhibited significantly delayed N100 latencies in the noise listening condition relative to the quiet listening condition (Non-Contact: Δ = 63 ms, *p* < .001; Contact: Δ = 57 ms, *p* < .001), with no group differences within either condition (Quiet: *p* = .255; Noise: *p* = .615). Figure 2 illustrates preserved cortical timing across groups despite amplitude reductions in contact athletes, indicating that sub-concussive head impacts selectively impair auditory response strength.

**Figure 2.**
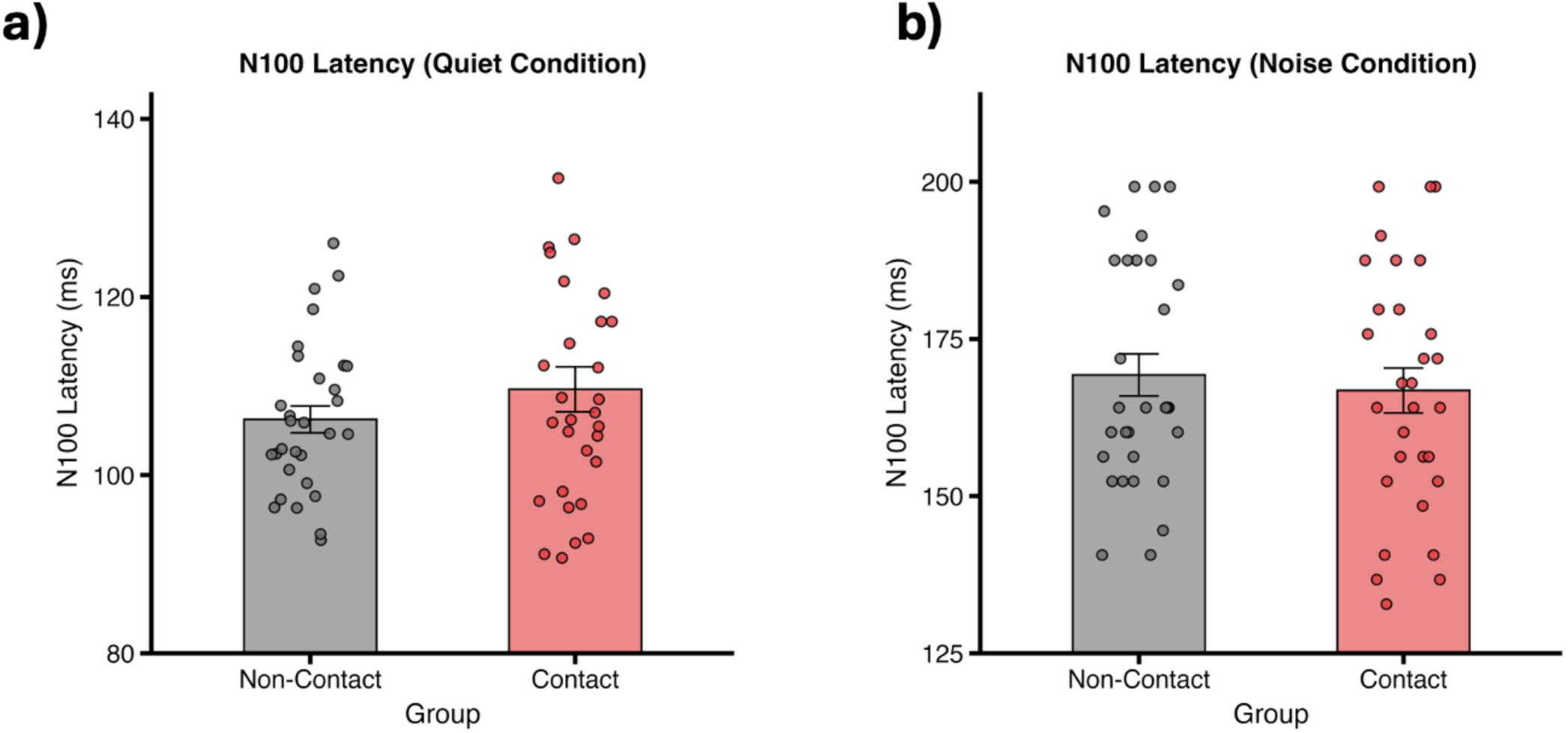
Cortical response timing is preserved despite group differences in response amplitude. (a) N100 peak latency in the quiet condition does not differ between contact and non-contact athletes, indicating comparable timing of early cortical sound encoding across groups. (b) Similarly, N100 latency in the speech-in-noise condition is statistically equivalent between groups, reinforcing that sub-concussive head impacts primarily affect response amplitude rather than neural timing.

### Sub-concussive Head Impacts Do Not Affect Subcortical Auditory Function

To examine whether repetitive sub-concussive head impacts affect subcortical encoding of the speech fundamental frequency (F0), a 2×2 mixed ANOVA assessed the effects of Group (Contact vs Non-Contact) and Condition (Quiet vs Noise) on F0 periodicity strength. Figure 3 illustrates the distribution of F0 encoding across groups and listening conditions, revealing no clear group differences. The effect of Group was not significant (F(1, 54) = 0.51, *p* = .476, η_p_²=.009), nor was the effect of Condition (F(1, 54) = 1.30 *p* = .258, η_p_² =.024) nor the Group × Condition interaction (F(1, 54) = 0.08, *p* = .776, η_p_² =.002).

**Figure 3.**
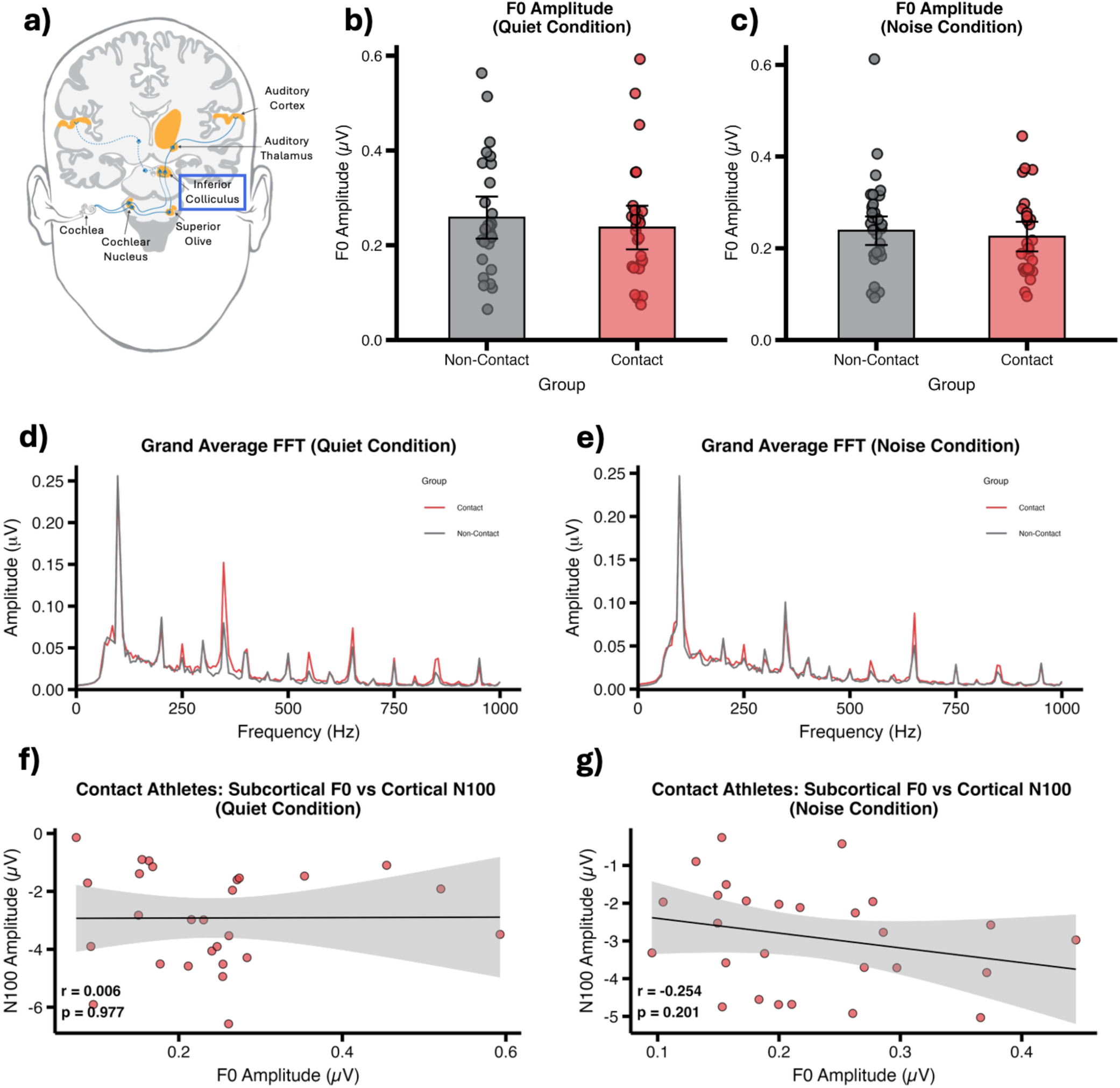
Repetitive head impacts do not disrupt subcortical periodicity encoding of speech. (a) *Schematic of the human auditory pathway highlighting subcortical regions thought to contribute to F0 periodicity encoding, particularly the inferior colliculus (adapted from Swords et al., 2018). (b) F0 amplitude in quiet shows no significant difference between contact and non-contact athletes. (c) F0 amplitude in speech-in-noise likewise reveals equivalent subcortical responses across groups. (d) Grand-average FFT spectra in the quiet condition reveal overlapping F0 peak amplitudes between contact and non-contact athletes, indicating preserved subcortical periodicity encoding. (e) In the speech-in-noise condition, FFT spectra similarly show no group differences in F0 amplitude, confirming intact subcortical responses under degraded listening conditions. (f) No association exists between individual F0 and cortical N100 amplitudes in the quiet condition. (g) A similarly weak, non-significant relationship between F0 and N100 amplitudes is observed in speech-in-noise, suggesting that cortical vulnerability to repetitive sub-concussive impacts occurs independently of subcortical periodicity encoding*.

Within the contact group, F0 amplitudes showed no correlation with cortical N100 amplitudes in either quiet (r = 0.006, *p* = .977, Fig. 3f) or noise conditions (r = -0.254, *p* = .201, Fig. 3g). These null subcortical findings, combined with robust cortical N100 amplitude reductions, provide evidence for a selective cortical signature of sub-concussive head impact effects, potentially highlighting cortical auditory processing as a sensitive biomarker of repetitive head impact exposure.

### Subcortical response timing preserved across listening conditions

To examine effects on subcortical response timing, a 2×2 mixed ANOVA assessed Group (Contact vs Non-contact) and Condition (Quiet vs Noise) on stimulus-to-response FFR latency. There was no significant main effect of Group (F(1, 54) = 0.62, *p* = 0.434, η_p_² = 0.011), confirming comparable subcortical timing across athlete groups. A significant main effect of Condition showed longer latencies in Noise versus Quiet (F(1, 54) = 10.51, *p* = 0.002, η_p_² = 0.163), with a non-significant Group × Condition interaction (F(1, 54) = 3.85, *p* = 0.055, η_p_² = 0.066).

Bonferroni-adjusted pairwise comparisons revealed no group differences in either condition (quiet: *p* = 0.70; noise *p* = 0.14). Within-group effects showed delayed latencies from quiet to noise in non-contact athletes (*p* < .001) but not contact athletes (*p* = 0.38). Click-evoked Auditory Brainstem Response (ABR) Wave V latency was also equivalent between groups (t(52) = -1.06, *p* = 0.29; non-contact *M* = 7.12ms vs contact *M* = 7.29ms). Figure 4 illustrates preserved sub-cortical timing across multiple measures despite cortical amplitude deficits in contact athletes.

**Figure 4.**
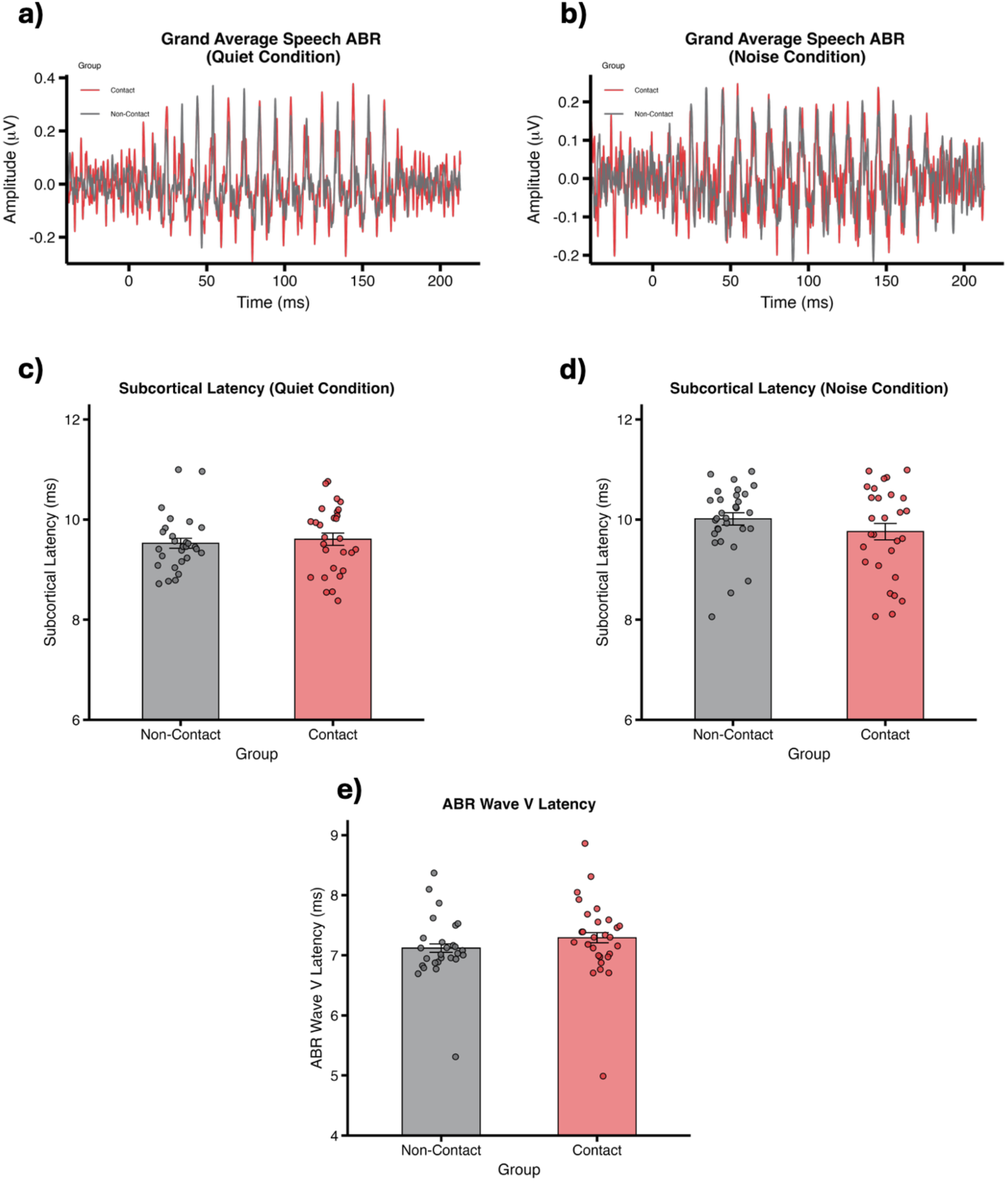
Subcortical response timing remains preserved across all neural levels despite sub-concussive head impacts. (a) Grand-average speech FFRs (/da/ stimulus) in the quiet condition show overlapping neural responses, meaning that there are no delays in processing speech sounds at a subcortical level between contact and non-contact athletes. (b) Speech FFRs in the speech-in-noise condition similarly reveal no group differences in morphology or timing. (b)Cross-correlation lag analysis confirms equivalent subcortical timing to speech stimuli in quiet between contact and non-contact athletes. (d) Subcortical timing to speech-in-noise is likewise statistically indistinguishable across groups. (e) Click-evoked ABR Wave V latency, a standard check for basic subcortical conduction delays, shows no group differences, verifying preserved early brainstem timing for both the contact and non-contact athletes.

### Auditory, Behavioural and Vestibular Measures Remain Unaffected by Sub-Concussive Head Impacts

Descriptive statistics for secondary auditory, behavioural and physiological measures are provided in Supplementary Table S1, confirming no significant group differences except for PTA thresholds, and SSQ12 Speech Subscale scores reported above. Postural sway metrics from PROTXX showed no significant group differences in eyes-open sway power t(33) = -1.70, *p* =.99, *d* = -0.44 (Fig. 5a) or eyes-closed sway power t(31) = -1.82, *p* = 0.79, *d* = -0.47 (Fig. 5b), despite a non-significant trend towards greater sway power in contact athletes with eyes closed. Figure 5 illustrates preserved postural stability across groups, providing further evidence that cortical auditory dysfunction represents the primary neural signature of repetitive sub-concussive head impact exposure.

**Figure 5.**
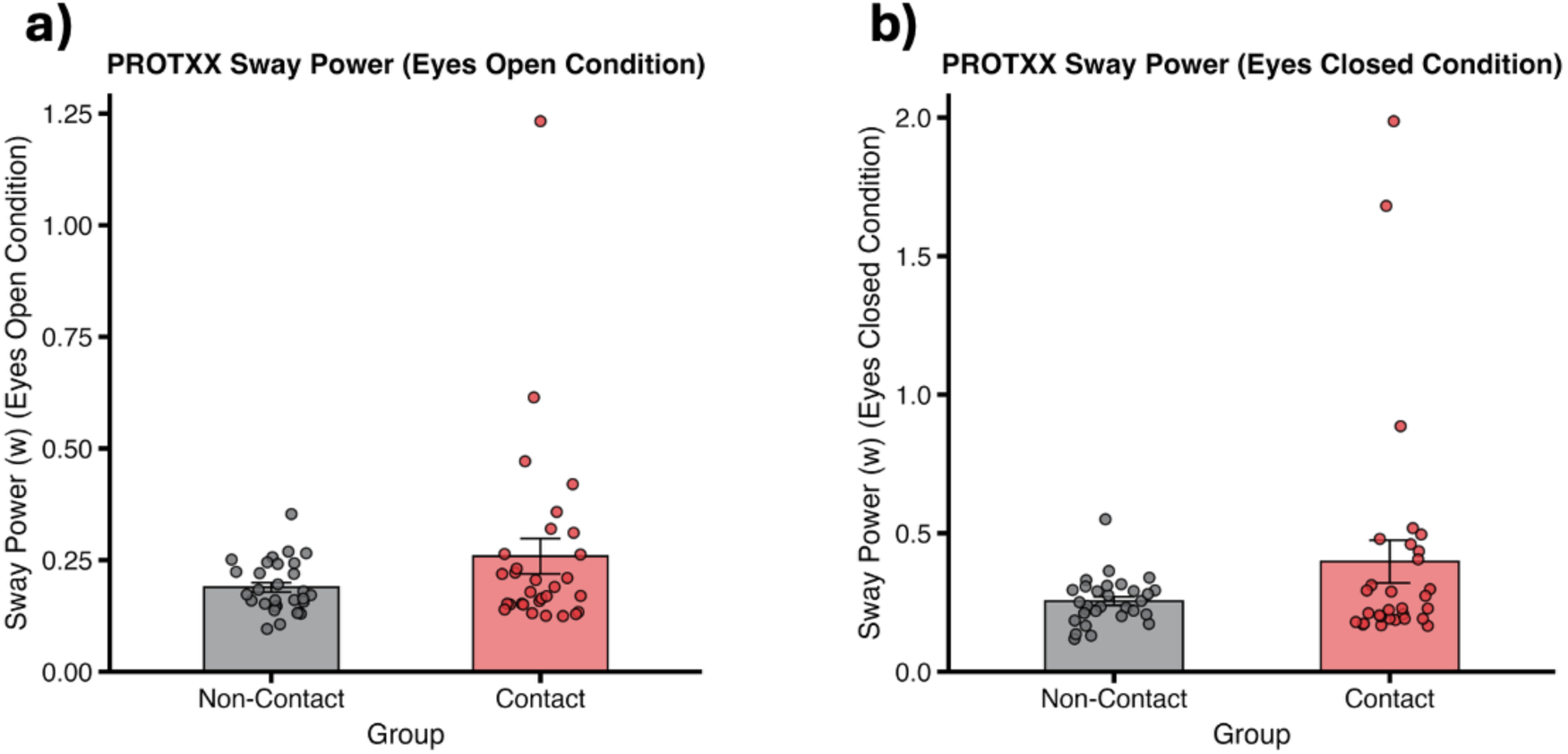
Postural stability is preserved in contact-sport athletes. (a) Sway power in the eyes-open condition shows no significant differences between contact and non-contact athletes. (b) In the eyes-closed condition, sway power reveals a non-significant trend toward greater sway in contact athletes, yet remains statistically equivalent across groups, indicating preserved vestibular and proprioceptive balance control.

## Discussion

Our findings uncover a novel and critical auditory cortical biomarker of repetitive sub-concussive head impacts in adult contact sport athletes, characterised by a pronounced reduction in N100 amplitude. Data indicate a significant disruption in early auditory cortical processing, which may represent one of the first neurophysiological markers of cumulative sub-concussive head impacts in this population of young adult contact sports athletes. Importantly, this study is the first to compare both subcortical and cortical responses within athletes, observing a specific cortical vulnerability to repetitive sub-concussive head impacts that is not evidence in subcortical responses. This finding furthers understanding of the neural mechanisms affected by repetitive sub-concussive head impacts, and how sub-concussive head impacts differentially affect central auditory processing. These insights will support the development of auditory neural biomarkers that could transform monitoring and intervention techniques for brain health in athletes who are exposed to repetitive sub-concussive head impacts.

The significant main effect of a reduction in the N100 amplitude, is potentially a result of subtle microstructural or functional changes at a cortical level following exposure to repetitive sub-concussive head impacts. This reduction in amplitude is likely to arise from a loss of synchronised firing and/or a decrease in the number of neurons processing auditory stimuli^[28]^. These interpretations are supported by recent pathological evidence^[18]^, which demonstrate significant neuron loss and inflammation in the cortical sulcal depths of young athletes exposed to repetitive head impacts. A dominant neural generator of the N100 is the primary auditory cortex^[29]^. Given the anatomical location of primary auditory cortex within the lateral sulcus, it is plausible that the cellular damage observed by Butler and colleagues^[18]^ could underlie the neural dysfunction revealed in the current study via the N100 response. Taken together, these data suggest a compelling narrative as to how the accumulation of repetitive sub-concussive head impacts may compromise cortical integrity; an effect that does not appear to be attributed to peripheral hearing loss, and is not mirrored by the FFR measures, although the latter may not be sensitive to all forms of subcortical pathology.

Several recent studies have investigated how repeated sub-concussive and concussive head impacts influence the auditory N100 amplitude, extending understanding of neural markers sensitive to cumulative head impacts. Fickling et al. (2019) reported an increased auditory N100 amplitude in Canadian junior-level ice hockey players, who had an average age of 18.49 ± 1.03 years, following medically diagnosed concussions, a finding that contrasts with the expected reduction in N100 amplitude but may indicate altered inhibitory/excitatory neural balance post-injury^23^. In the same study, no significant N100 amplitude differences were observed related to sub-concussive head impact exposure. However, a subsequent study by Fickling et al. (2021) also with junior ice hockey players, aged 13.63 ± 0.62 years, but focusing on pre- to post-season changes reflective of cumulative sub-concussive head impact, found a significant decrease in N100 amplitude using a passive listening paradigm^[24]^. Further extending this work to youth American football players, aged 12.89 ± 0.35 years, Fickling et al., (2022) found that while N100 amplitude did not significantly decrease across a season, it was a significant predictor of repetitive sub-concussive head impacts, underscoring its utility as a sensitive biomarker for repetitive sub-concussive exposure even in the absence of overt group-level change^[30]^.

Further research has supported these findings, Canadian high school athletes, aged between 15 – 17 years old, who engaged in contact sports with exposure to repetitive head impacts also had a significant reduction in auditory N100 amplitude responses^[31]^. These electrophysiological changes are further supported by additional data from adult populations with medically diagnosed concussive head impacts and repetitive sub-concussive head impact exposure. For example, D’Arcy and colleagues^[45]^ found that athletes with a history of concussion had reduced N100 amplitudes during active cognitive auditory tasks, compared to the no-concussion group, while another study Munce and colleagues found that mixed martial arts athletes exposed to repetitive sub-concussive head impacts had reduced auditory N100 (alongside reduced P300 and N400) amplitudes, compared to matched controls^[33]^. Taken together, this growing body of evidence indicates that in a variety of athlete populations and experimental designs, reduced cortical auditory responses, particularly the N100 amplitude, are a sensitive indicator of concussive and cumulative sub-concussive head impacts, although there are some variations, particularly in younger athletes and under different task demands. The present study adds to this work as it indicates that findings of cortical dysfunction are likely not inherited from earlier subcortical generators, nor related to changes to peripheral hearing sensitivity. Our observation of N100 amplitude reduction in young adult contact sport athletes during passive listening adds important confirmation of central auditory processing dysfunction and advances the potential utility of N100 responses as an objective biomarker for brain health monitoring in at-risk populations exposed to repetitive sub-concussive head impacts.

The present study did not find any significant differences in subcortical auditory processing, measured using the F0 amplitude of the FFR, between contact and non-contact athletes in either quiet or noise conditions. In contrast, previous evidence has demonstrated subcortical auditory processing deficits following both concussive and repetitive sub-concussive head impacts^[34,35]^. This work has evidenced significant reductions in FFR measures of pitch and phonetic encoding^[25]^, which persist even after symptoms resolve and extend to younger populations, including children who have sustained a single concussion^[34]^. These long-lasting subcortical auditory effects indicate persistent neural impact of concussion and sub-concussion at earlier neural generators in the auditory pathway. However, replicating consistent subcortical auditory processing differences has proven challenging in other cohorts. For example, Van der Werff and Rieger^[22]^ found no significant differences in either latency or amplitude of brainstem responses to speech and click stimuli between individuals with mild traumatic brain injury (mTBI) and matched controls. Similarly, this study did not find subcortical F0 amplitude differences between contact and non-contact athletes.

The present study used a controlled cohort of male and female athletes aged 18 to 27 years involved in team- and invasion-based contact sports (e.g., rugby and football) and age-, sex-, height-, body mass- and BMI-matched non-contact athletes also from team- and invasion-based non-contact sports (e.g., netball and handball), yet did not observe any subcortical differences. Several factors may account for discrepancies with previous work reporting subcortical alterations. For example, one large-scale study done by Kraus and colleagues included athletes from multiple sporting disciplines with varying degrees of contact and head impact exposure, whereas the current study focused specifically on athletes involved in team-and invasion-based contact sports, applying rigorous criteria to define both contact and non-contact groups^[36]^. This targeted approach likely reduced variability within groups resulting from differences in sport type and frequency of repetitive sub-concussive head impacts, enabling a more controlled and internally consistent comparison. Moreover, the current study implemented rigorous screening procedures, including controlling for sex hormone variability among female participants, a known modulator of auditory neural responses^[37]^ which was not detailed in Kraus et al.’s research. In addition, the present study collected self-reported data on concussion history and ensured participants were cleared of concussion for at least one month before testing. This exclusion criterion is stricter than Kraus and colleagues who retained athletes with recent concussions but controlled for the number of past concussive head impacts as a covariate in their ANOVA^[49]^. By physically removing recent concussion effects rather than statistically adjusting for them, the current study more effectively minimised the influence of acute concussion effects. This cleaner approach enhances the accuracy of FFR-F0 amplitude as a marker for cumulative or longer-term brain alterations linked to repetitive sub-concussive head impact exposure is improved by this strict control, which lowers the possibility of confounding from recent injury. Thus, our null subcortical findings, obtained under stricter exclusion criteria, suggest that previously reported FFR differences may have reflected residual effects of recent concussions rather than a reliable biomarker of repetitive sub-concussive head impacts.

Importantly, exploratory analyses from the present study revealed no significant correlations between cortical N100 amplitude and subcortical F0 amplitude within the contact athlete group in either quiet or noise conditions. Additionally, contact athlete’s PTA thresholds for both ears, or their SSQ12 Speech subscale scores did not correlate significantly with either cortical or subcortical auditory measures. These findings suggest that the cortical N100 reductions observed in contact athletes were not accompanied by parallel changes in subcortical F0 responses or peripheral and behavioural hearing sensitivity. This dissociation supports the claim that the observed cortical N100 reductions represent a distinct neural signature of repetitive sub-concussive head impacts, rather than being a downstream consequence of subcortical dysfunction or changes to peripheral hearing sensitivity.

These findings have important implications that extend beyond the identification of potential biomarkers for athlete health. They also point to a commonly overlooked group of athletes who may be suffering from central auditory processing difficulties. These challenges often present as subtle auditory complaints that increase cognitive effort during everyday listening. In this way, according to the cognitive load hypothesis, central auditory processing deficits and peripheral hearing loss contribute to cognitive decline and elevate dementia risk^[38]^. In those athletes exposed to repeated sub-concussive head impacts, such central processing issues may further contribute to the broader neurodegenerative processes involved in conditions characterised by tau pathology^[39]^. These central auditory processing deficits therefore warrant recognition when considering clinical monitoring and intervention strategies for this group, as they may impact long-term brain health and cognitive function.

## Clinical and Monitoring Relevance

The exposure-related reduction in cortical N100 amplitude identified in this study has important clinical and monitoring implications. As an objective neurophysiological marker of repetitive sub-concussive head impacts, N100 amplitude measurement may facilitate early detection of subtle brain changes before overt symptoms emerge, enabling proactive monitoring and intervention in contact sport athletes. Given that these electrophysiological measures are non-invasive and can be obtained during passive listening, they are well-suited for integration as part of regular screening sessions between and during seasons of play for at-risk athletes.

## Limitations and Future Directions

A key strength of this study is the use of a rigorously defined sample of athletes primarily participating in team- and invasion-based contact sports, enhancing internal validity. However, while groups were age-, sex- height-, body mass-, BMI-matched, other lifestyle factors that could potential differ between contact and non-contact athletes, such as alcohol consumption, recreational drug use and loud music exposure, were not recorded and could represent unmeasured confounds^[40]^. Concussion history was captured via self-report, and participants with a diagnosed concussion within the previous month were excluded. However, there is still a possibility of unreported or unrecognised concussive events within the sample, due to the challenges around diagnosing concussion, which could have impacted the observed outcomes. Additionally, precise quantification of sub-concussive head impact exposure was not available, preventing direct correlation of the observed auditory changes with the amount or severity of impacts. Future studies should incorporate wearable accelerometers, or video-assisted impact monitoring to objectively capture head impact exposure to investigate dose-response relationships. Longitudinal designs to track neural changes over multiple seasons or years of play would help understand progression of exposure-related effects. Furthermore, exploration of auditory cortical and subcortical processing in retired athletes with extended histories of contact sport exposure may provide critical insights into whether the cortical changes observed during active participation maintain, reduce, or increase over time. Such research could help distinguish transient versus chronic neural effects and inform our understanding of potential long-term consequences of repetitive sub-concussive impacts. Finally, future studies expanding to include auditory behavioural assessments will strengthen the ecological validity by linking neural disruptions to functional impairments, and including a comprehensive lifestyle assessment and a broader range of ages will enhance understanding of exposure-outcome relationships across diverse athlete populations.

## Conclusion

This study is a critical step towards understanding neural vulnerability to repetitive sub-concussive head impacts. By identifying a selective cortical processing difference in young adult contact sport athletes, we provide evidence of a sensitive and objective EEG-based biomarker of sub-concussive exposure. The absence of subcortical changes suggests that cortical alterations are not inherited from lower auditory structures, indicating distinct cortical susceptibility. While this biomarker arises from auditory neural activity, its significance extends beyond hearing: it offers a practical, non-invasive means to detect early brain changes that currently can only be confirmed post-mortem through neuropathological examination. These findings highlight the potential of electrophysiological markers to transform the early detection, monitoring, and management of repetitive head impact exposure, supporting athlete health, risk assessment, and safety in contact sports

## Materials and Methods

### Ethics and Regulatory Approvals

Ethical approval was obtained from the Faculty of Science and Technology Research Ethics Committee at Lancaster University FST-2024-4150-RECR-3 before any data collection commenced. All experimentation conformed to the *Declaration of Helsinki* (7^th^ edition, 2013). Informed consent was obtained from all participants prior to any study procedures, following completion of a self-reported screening questionnaire administered via Qualtrics and before attendance of the lab testing session The study was pre-registered on the Open Science Framework (OSF) prior to data collection (https://osf.io/e59sk).

### Participants

Participants were recruited from Lancaster University and local sports clubs. Of the 211 individuals who expressed interest, 60 met the eligibility criteria and completed the study. A detailed overview of participant screening and exclusion is provided in the CONSORT flow diagram (Supplementary Figure S1). The final sample comprised 30 contact-sport athletes (12 female, 18 male; aged 18–26 years, *M* = 20.3, *SD* = 1.9) and 30 non-contact sport athletes (17 female, 13 male; aged 18–27 years, *M* = 20.6, *SD* = 2.3), with no significant between-group differences in age, height, body mass, body mass index (BMI) or sex distribution (Table 1).

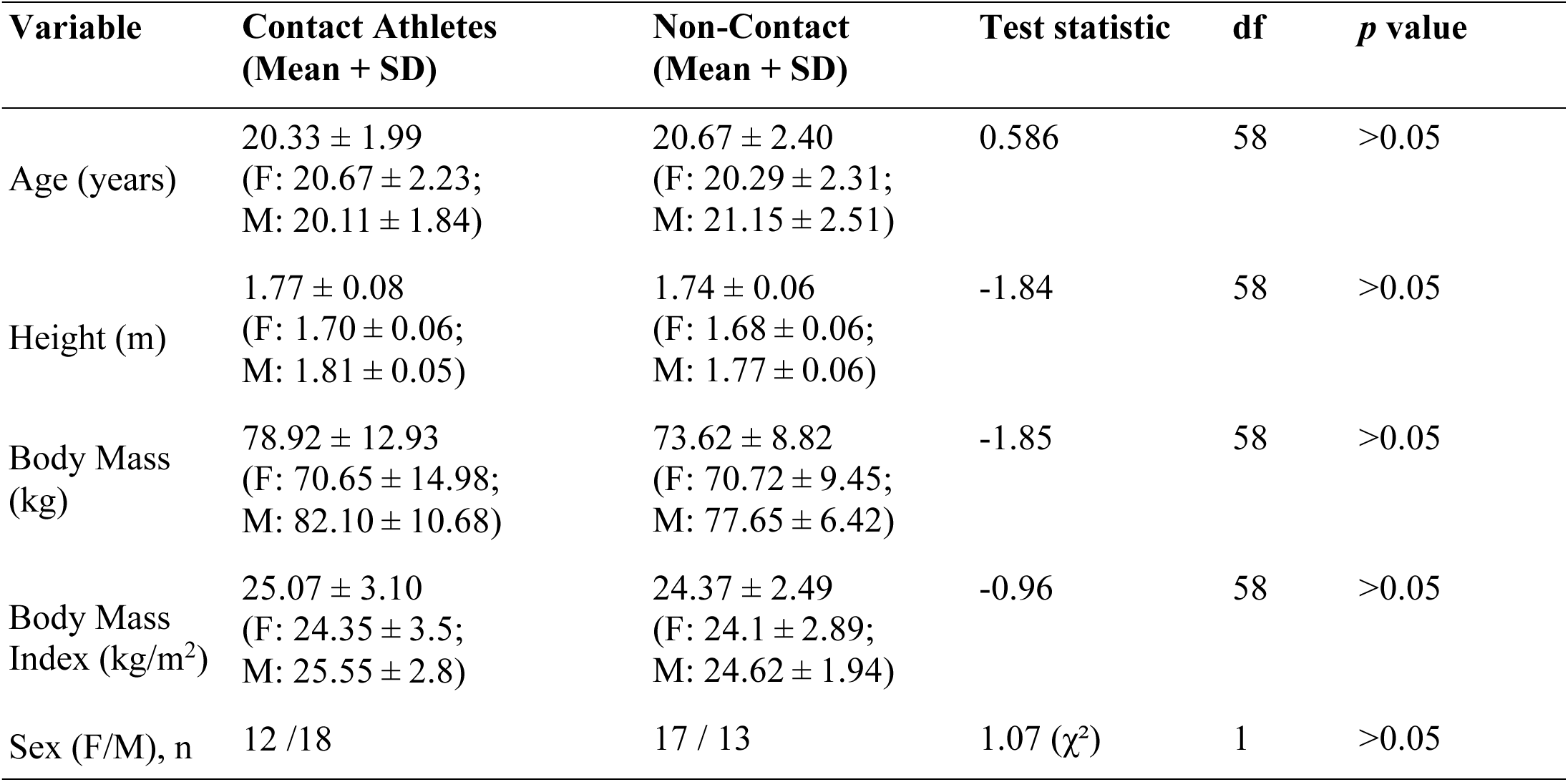

| Variable | Contact Athletes<br>(Mean + SD) | Non-Contact<br>(Mean + SD) | Test statistic | df | *p* value |
| --- | --- | --- | --- | --- | --- |
| Age (years) | 20.33 ± 1.99<br>(F: 20.67 ± 2.23;<br>M: 20.11 ± 1.84) | 20.67 ± 2.40<br>(F: 20.29 ± 2.31;<br>M: 21.15 ± 2.51) | 0.586 | 58 | >0.05 |
| Height (m) | 1.77 ± 0.08<br>(F: 1.70 ± 0.06;<br>M: 1.81 ± 0.05) | 1.74 ± 0.06<br>(F: 1.68 ± 0.06;<br>M: 1.77 ± 0.06) | -1.84 | 58 | >0.05 |
| Body Mass<br>(kg) | 78.92 ± 12.93<br>(F: 70.65 ± 14.98;<br>M: 82.10 ± 10.68) | 73.62 ± 8.82<br>(F: 70.72 ± 9.45;<br>M: 77.65 ± 6.42) | -1.85 | 58 | >0.05 |
| Body Mass<br>Index (kg/m<sup>2</sup>) | 25.07 ± 3.10<br>(F: 24.35 ± 3.5;<br>M: 25.55 ± 2.8) | 24.37 ± 2.49<br>(F: 24.1 ± 2.89;<br>M: 24.62 ± 1.94) | -0.96 | 58 | >0.05 |
| Sex (F/M), n | 12 / 18 | 17 / 13 | 1.07 ($\chi^2$) | 1 | >0.05 |

Participants were eligible if aged between 18-30 years, monolingual native British English speakers^[41]^, and competing at Tier 2 level in invasion-based contact or non-contact sports^[42]^, defined as team games in which players invade an opponent’s territory to score while defending their own^[43]^. Female participants eumenorrheic status without menstrual/reproductive conditions, with testing during days 1-5 of the menstrual cycle to control for hormonal effects of auditory sensitivity^[37]^. Exclusions included extensive musical training, history of auditory processing disorder, childhood hearing loss, or neurological/neurobiological conditions, as these may confound auditory processing mechanisms^[44,45]^. A breakdown of sports represented is provided in Supplementary Table S2.

### Sample Size Calculations

An a-priori calculation was performed using G*Power 3.1 (version 3.1.9.6)^[46]^ for a 2 x 2 mixed model analysis of variance (ANOVA). Calculation parameters were a medium effect size (f = 0.25) with a statistical power of 80% and an alpha value of 0.0125, which was corrected for family-wise error rate across the four hypotheses (0.05/4). A medium effect size was selected to reflect the range of effect magnitudes previously observed across auditory processing levels in contact sport athletes and to balance sensitivity with practical feasibility^[31,36]^. The sample size calculation indicated that a total sample of 48 participants would be required, with an equal distribution in each group (contact sport athletes, n = 24, non-contact sport athletes, n = 24). To account for any potential data loss or dropouts, we aimed to collect 30 participants in each group, making a total of 60 participants. All 60 participants provided complete datasets, with a small number of data points later excluded due to quality issues (see Statistical Analysis for details)

### Experimental Design

This research study implemented a 2 (Group: contact vs non-contact sport athletes) x 2 (Condition: Quiet listening condition vs speech-in-noise listening condition) mixed model design, with group as the between-subjects factor and condition as the within-subjects factor.

The experiment was completed in a single session, using two types of EEG responses: the (FFR) and the N100. Participants completed both response types in two listening conditions: one where the stimuli were presented in quiet and one in a speech-in-noise environment. The order in which participants completed the FFR and N100 recordings, as well as the order of the two listening conditions was counterbalanced across participants to control for order effects.

## Demographic Variables

### Speech, Spatial and Quality of Hearing Questionnaire (SSQ12)

Participants completed the 12-item Speech, Spatial and Qualities of Hearing Scale (SSQ12), a validated short-form questionnaire derived from the original 49-item SSQ^[48]^. The SSQ12 assesses subjective hearing abilities in three domains (1) speech perception in complex listening environments, (2) spatial hearing (e.g., localisation and movement detection), and (3) auditory qualities (e.g., sound segregation and listening effort). Participants rated their hearing ability in different acoustic scenarios using a sliding scale from 0 to 10 (0 = not at all, 10 = perfectly) on Qualtrics, capturing self-reported experiences of auditory function.

### PROTXX

Participants completed a postural sway assessment using PROTXX (PROTXX Inc, Menlo Park, CA, USA), a wireless physiological vibration acceleration (phybrata) sensor^[49]^. Participants were first asked if they had any skin sensitivities or allergies to adhesive materials, such as plasters. If sensitivities were identified, PROTXX inertial measurement unit (IMU) was secured using a headband, otherwise, a disposable medical adhesive patch was used to place the sensor on the right mastoid. Participants stood upright with feet hip-width apart, arms by their sides, and maintained a fixed gaze on a target 3 metres away. They were instructed to remain still, avoid talking, chewing gum, head movements, or fidgeting during the test. Using the protxxclinic smartphone app (Version 1.0 build 13), connected via Bluetooth, two 60-second trials were conducted, (1) eyes open and (2) eyes closed. The app provided audible countdowns and a tone ten seconds before each trial ended. A researcher supervised the participant, especially during eyes-closed trials, to ensure safety. The PROTXX device recorded acceleration along three axes (front-back, vertical, left-right) at 100 Hz, producing 12,000 samples per axis per trial. Data were filtered with a 0.04 Hz high-pass filter to remove drift and gravitational bias. The software calculated sway magnitude for eyes open and eyes closed, their ratio, and average power, the latter considered a more objective measure of balance impairment and concussion severity than single conditions alone^[49]^.

## Hearing Assessment

### Otoscopic Examination

A visual inspection of the ear canal and eardrum was performed using a handheld otoscope. This was to check for any visible damage to the eardrum (e.g., holes, scarring) or signs of infection (e.g., redness, fluid). Participants were excluded if earwax buildup or blockages were observed, as these could interfere with accurate hearing measurements. No participants were excluded based on this examination.

### Pure-Tone Audiometry

Hearing thresholds were measured bilaterally using air-conduction pure-tone audiometry across frequencies ranging from 250 Hz to 8000 Hz (250, 500, 1000, 2000, 4000, and 8000 Hz), following British Society of Audiology (BSA) guidelines^[50]^ using a calibrated Amplivox Model 270 Diagnostic audiometer (Amplivox LTD, UK).

### Auditory Brainstem Response (ABR)

Click-evoked ABRs were recorded to confirm normal brainstem function. Wave V peak latencies fell within expected normative ranges for all participants (*M*= 7.21 ms, *SD* = 0.61), indicating intact auditory nerve lower brainstem transmission.

### EEG

Both cortical and subcortical EEG data were acquired using the BioSemi ActiveTwo EEG system (BioSemi, Amsterdam). For cortical responses, a 32-channel scalp montage was recorded using Speed Mode 4 with a sampling rate of 2,048 Hz. Signals were referenced to the left earlobe, and no ocular or facial electrodes were recorded. For subcortical recordings, Speed Mode 9 was used with a sampling rate of 16,384 Hz. A vertical electrode montage was employed, with the active electrode placed at Cz, referenced to the left earlobe, and two ground electrodes positioned on the forehead. Electrode sites were prepared using alcohol wipes to reduce impedance, ensuring values were below 5 kΩ. The Cz electrode was determined using standard 10–20 system measurements, with the nasion and inion as anatomical landmarks.

### EEG Stimuli

Stimuli (click, speech, and multi-talker babble) were calibrated in dB SPL using a Brüel and Kjær type 2250 (Brüel and Kjær, Nærum, Denmark) sound-level meter and presented binaurally via RadioEar IP30 insert earphones (RadioEar, US). The trigger timing was calibrated and aligned with the sound output using the Black Box ToolKit v2 (The Black Box ToolKit Ltd., 2016).

### Click stimulus

A brief click stimulus (100 μs duration) was used to elicit the ABR. The clicks were presented at 70 dB nHL using alternating polarity (condensation and rarefaction) at a rate of 11.1 clicks per second (2200 total presentations: 1100 per polarity) to eliminate cochlear microphonic (CM) artefacts, which invert with polarity changes and cancel out in the averaged response. This approach ensures that the recorded ABR reflects neural activity rather than CM contributions^[51]^.

### Speech stimulus: 170ms /da/

A 170 ms speech syllable /da/ was synthesised using a Klatt synthesizer^[52]^ at a 20-kHz sampling rate and presented at 70 dB SPL. This stimulus, originally developed by Kraus and colleagues at the Auditory Neuroscience Laboratory, Northwestern University, consists of a brief onset phase, a formant transition period, and a steady-state period, see Skoe and Kraus^[53]^ for further details.

### Noise stimulus

For the noise condition, a six-talker babble masker (three female, three male) was created using randomly selected recordings from the Clarity Speech Corpus^[54]^. Babble files were generated to ensure equal talker representation and a dynamic, non-repetitive background, and were presented at 60 dB SPL during stimulus presentation to achieve a +10 dB signal-to-noise ratio (SNR). Further details of the babble construction and MATLAB scripts used are available in Supplementary Material (see S1 file).

## EEG procedures

Stimulus presentation was controlled using Neurobehavioral Systems, Presentation software (Neurobehavioral Systems, Inc., Albany CA). Throughout testing, participants watched a muted nature documentary with subtitles removed to minimise cognitive and language-related distractions^[55]^. To facilitate a passive-listening state, participants watched a nature documentary during recording. As the auditory stimulus was presented binaurally, the documentary was muted, and subtitles were removed to minimise auditory and linguistic interference, as reading and language-related tasks can activate auditory pathways, potentially modulating neural outcomes^[56]^. EEG data were collected separately for FFRs and N100 responses using the same stimulus, but different acquisition settings optimised for the temporal and physiological characteristics of each response type.

For the cortical recordings, a longer interstimulus interval (ISI) of 863 ms was used to avoid overlap of slow cortical potentials, resulting in a total trial length of 1,033 ms. Each condition included 200 trials, corresponding to a recording time of approximately 3 minutes and 26 seconds of recording per condition.

For the subcortical recordings, the /da/ stimulus was presented with alternating polarity, a method used to suppress cochlear and stimulus artefacts while enhancing the phase-locked neural signal. Each condition consisted of 2,200 stimulus presentations (1,100 per polarity) with an ISI of 83 ms, resulting in a total trial duration of 253 ms and a recording time of approximately 9 minutes and 18 seconds per condition.

## EEG Pre-Processing

EEG data were pre-processed offline in MATLAB R2025a using EEGLAB version 2024.0^[57]^. Two separate preprocessing pipelines were applied for subcortical and cortical EEG recordings due to the distinct temporal and neural characteristics of the responses of interest^[58]^.

### Subcortical EEG Pre-Processing

Subcortical EEG data were retained at the original 16,384 Hz sampling rate to preserve high-frequency information critical for brainstem analyses. Data were first re-referenced to the left earlobe. A bandpass filter (FIR, zero-phase) was applied using pop_eegfiltnew to isolate frequencies between 70 Hz (high-pass) and 2000 Hz (low-pass) settings optimised for capturing phase-locked FFRs. The continuous recordings were then segmented into epochs ranging from - 40 ms to 213 ms relative to stimulus onset. Trials exceeding ±75 μV were automatically rejected. Independent component analysis (ICA) and additional ERP Lab-based cleaning steps were intentionally omitted, as the short epoch duration and highly phase-locked nature of FFRs make these techniques potentially disruptive to the signal. Baseline-correction was applied to all retained epochs.

### Cortical EEG Pre-Processing

Cortical EEG data were downsampled to 256 Hz after import to reduce computational demand, a deviation from the original preregistration. The data were re-referenced to the left earlobe and then bandpass filtered using EEGLAB’s pop_eegfiltnew function, with cut-offs set at 1 Hz (high-pass) and 20 Hz (low-pass) to isolate cortical evoked potentials and suppress noise. ICA was performed using the Infomax algorithm (pop_runica) to identify and remove physiological artefacts such as blink and muscle activity. ICA components were automatically labelled using ICLabel and flagged based on standard threshold criteria.

Following ICA, cleaned data were segmented into epochs from -100 ms to 400 ms relative to stimulus onset. Epochs with amplitudes exceeding ±50 μV were excluded. Baseline correction was applied using the interval from -100 to 0 ms, helping to remove residual low-frequency drift and align responses across trials.

## Exploratory EEG Latency Measures

Latency measures were explored as supplementary outcomes to quantify the temporal dynamics of the auditory neural response at both subcortical and cortical levels. Latency was quantified as follows:

### Cortical (N100)

Cortical latency using the latency of the N100 component, which occurs approximately 100 ms after stimulus onset. EEG data were recorded at electrode Cz, and latency extraction was performed on cleaned, baseline-corrected epochs using a combination of automated slope detection and visual validation. Specifically, the most negative deflection within a post-stimulus window of 80 to 180 ms was identified for each participant, corresponding to the N100 peak.

### Subcortical Latency (Stimulus-to-Response)

Subcortical latency was quantified using a stimulus-to-response cross-correlation (XCorr, MATLAB^[59]^) technique, which quantifies the temporal lag between the presented auditory stimulus and the phase-locked subcortical response. For each participant, the EEG response at Cz was averaged across artefact-free sweeps and compared with the original 170-ms /da/ stimulus waveform. To align the neural and stimulus signals, the stimulus was resampled to the EEG sampling rate (16,384 Hz) and both waveforms were bandpass filtered (70–2000 Hz) using a second-order zero-phase Butterworth filter. The EEG response was then segmented from 40 to 210 ms, corresponding to align with the acoustic duration of the 170-ms /da/ stimulus, in order to isolate the speech-evoked portion of the subcortical response. Cross-correlation was performed over a constrained lag window of 8 to 11 ms, which accounts for both neural transmission time and the ∼1 ms acoustic delay introduced by insert earphones. The lag value (in milliseconds) at which maximum correlation occurred was extracted for each participant and interpreted as the subcortical stimulus-to-response latency.

### Statistical Analysis

All statistical analyses were conducted in RStudio (Version 2023.06.0+421). The following packages were used: readxl (data import), dplyr and purrr (data wrangling), car (assumption testing), afex (ANOVA models), emmeans (post-hoc-pairwise comparisons), broom (model tidying) and ggplot2 (visualisation).

Descriptive statistics (means, standard deviations, and ranges) were calculated for all primary and secondary variables. A small number of data points were excluded within the primary variables due to quality concerns, including noisy or poorly defined waveforms and implausibly high values suggesting artefact contamination. For the F0 amplitude analyses, inclusion rates were as follows: 96.7% of non-contact participants in the quiet condition, 100% in the speech-in-noise condition, 93.3% of contact participants in quiet, and 90.0% in speech-in-noise. For N100 amplitude, 96.7% of non-contact participants in quiet, 100% in speech-in-noise, 93.3% in the contact group for quiet, and 100% in speech-in-noise were included in the final analyses

Auditory response measures were performed in MATLAB (R2025a) using EEGLAB. The main cortical response measure was the N100 amplitude, extracted from electrode Cz within a post-stimulus latency window of 80–180 ms. Peak amplitudes and latencies were automatically detected using an automated slope-based peak detection. These values were then visually confirmed for accuracy on a participant-by-participant basis, ensuring robust and consistent identification of cortical auditory responses. To assess reliability of pitch encoding, the F0 amplitude was later extracted using fast Fourier transform (FFT) applied to averaged EEG responses from Cz, focusing on the amplitude within a 90–110 Hz band.

Two separate two-way mixed-model ANOVAs were conducted to test the primary hypotheses, one for each main dependent variable: (1) cortical N100 amplitude and (2) subcortical F0 amplitude from the FFR. Each ANOVA included Group (contact vs. non-contact athletes) and Condition (speech in quiet vs. noise) as independent variables. All ANOVA assumptions were checked prior to interpretation. Bonferroni correction adjusted the alpha level to *p* < .0125 to control for multiple comparisons across the four pre-specified hypotheses. Significant effects were further explored with Bonferroni-corrected post hoc pairwise comparisons, with a significance set at *p* < .025. these comparisons assessed between group differences within each condition and within-group differences between conditions.

## Acknowledgements

The authors gratefully acknowledge the Economic and Social Research Council (ESRC) for funding this project through an ESRC industrial CASE Studentship (UKRI Grant Code: ES/P000665/1; Project: 2700468a) with CASE partner Brainbox Ltd. We also thank the participating athletes and sports teams for their time and commitment to the study, and colleagues at Lancaster University and Brainbox Ltd. for technical assistance and data collection support.

## Data Availability Statement

Data and code will be made publicly available via the project listing on the Open Science Framework (https://osf.io/e59sk) following publication. Supplementary material, including additional analyses and methodological details, will be provided alongside the published article and hosted on the same project page.

