## Supplementary materials for "Distinct Neural Signatures of Auditory Processing in Contact versus Non-contact Sports Athletes"

**Supplementary Figure S1. CONSORT participant flow diagram**

*Flow of participants through the study, from initial interest (n=211) to final inclusion in analyses (n=60), showing exclusion criteria at each screening stage*

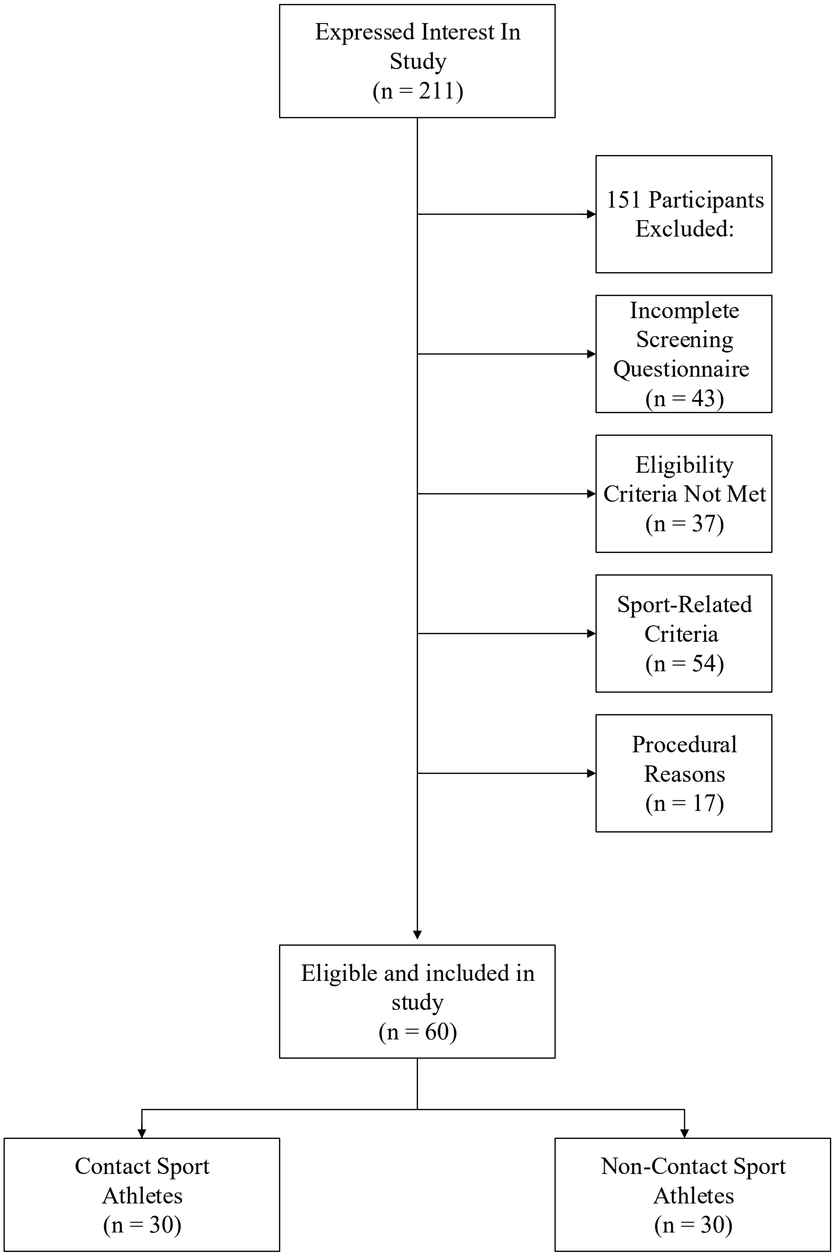

**Supplementary Table S1. Group differences in self-report, hearing, and postural sway measures**

*Means, standard deviations, and ranges for Contact and Non-Contact groups, with independent-samples t-tests (Welch) comparing groups on self-report (SSQ12), pure-tone average (PTA), PROTXX sway power, and click-evoked ABR Wave V latency. Significance is indicated at α = .05*

| Variable | Group | Mean | SD | Min | Max | t(df) | *p* | Sig? |
|---|---|---|---|---|---|---|---|---|
| SSQ12 Speech | Contact | 7.38 | 1.45 | 4.6 | 9.6 | 2.67 (53.97) | .010 | **Yes** |
| | Non-Contact | 8.3 | 1.1 | 6 | 10 | | | |
| SSQ12 Spatial | Contact | 7.89 | 1 | 10 | 6.34 | 0.08 (50.61) | .941 | No |
| | Non-Contact | 7.92 | 1.26 | 5 | 10 | | | |
| SSQ12 Hearing Quality | Contact | 6.37 | 1.6 | 3.5 | 9.5 | –1.04 (57.98) | .304 | No |
| | Non-Contact | 5.94 | 1.57 | 2.75 | 10 | | | |
| SSQ12 Overall | Contact | 7.17 | 1.22 | 4.08 | 9.17 | 0.88 (50.62) | .383 | No |
| | Non-Contact | 7.41 | 0.81 | 5.83 | 10 | | | |
| PTA Left Ear (dB) | Contact | 8.33 | 5.15 | 1 | 20 | –2.69 (53.24) | .010 | **Yes** |
| | Non-Contact | 5.2 | 3.7 | 0 | 13 | | | |
| PTA Right Ear (dB) | Contact | 7.77 | 5.22 | -3 | 17 | –2.15 (51.75) | .036 | **Yes** |
| | Non-Contact | 5.27 | 3.63 | -2 | 13 | | | |
| PTA Better Ear (dB) | Contact | 6.69 | 5.26 | -3 | 17 | -2.59 (48.36) | 0.01 | **Yes** |
| | Non-Contact | 3.77 | 3.26 | -2 | 12 | | | |
| PROTXX Postural Sway Power (Eyes Open) | Contact | 0.26 | 0.23 | 0.12 | 1.24 | –1.70 (33.13) | .099 | No |
| | Non-Contact | 0.19 | 0.06 | 0.1 | 0.35 | | | |
| PROTXX Postural Sway Power (Eyes Closed) | Contact | 0.4 | 0.42 | 0.17 | 2 | –1.82 (31.33) | .079 | No |
| | Non-Contact | 0.25 | 0.08 | 0.12 | 0.55 | | | |
| PROTXX Postural Sway Power (EC/EO Ratio) | Contact | 1.52 | 0.93 | 0.91 | 6.21 | –0.84 (34.95) | .408 | No |
| | Non-Contact | 1.37 | 0.3 | 0.92 | 2.39 | | | |
| Click ABR Wave V Latency (ms) | Contact | 7.29 | 0.66 | 5.00 | 8.85 | –1.06 (51.73) | .290 | No |
| | Non-Contact | 7.12 | 0.54 | 5.31 | 8.36 | | | |

**Supplementary Table S2**. **Sports represented by contact and non-contact athletes**

*Distribution of sports within contact (n=30) and non-contact (n=30) groups by sex.*

| **Sport** | **Contact** | | **Non-Contact** | |
|---|---|---|---|---|
| | Female | Male | Female | Male |
| Football | 6 | 5 | - | - |
| Rugby | 2 | 8 | - | - |
| Rugby Union | 4 | 3 | - | - |
| American Football | 0 | 1 | - | - |
| Lacrosse | 0 | 1 | - | - |
| **Contact total** | **12** | **18** | - | - |
| Netball | - | - | 14 | 0 |
| Hockey | - | - | 3 | 1 |
| Basketball | - | - | 0 | 5 |
| Korfball | - | - | 0 | 4 |
| Frisbee | - | - | 0 | 1 |
| Handball | - | - | 0 | 2 |
| **Non-contact total** | - | - | **17** | **13** |
